# Surveillance to Establish Elimination of Transmission and Freedom from Dog-mediated Rabies

**DOI:** 10.1101/096883

**Authors:** Katie Hampson, Bernadette Abela-Ridder, Kirstyn Brunker, S. Tamara M. Bucheli, Mary Carvalho, Eduardo Caldas, Joel Changalucha, Sarah Cleaveland, Jonathan Dushoff, Veronica Gutierrez, Anthony R Fooks, Karen Hotopp, Daniel T Haydon, Ahmed Lugelo, Kennedy Lushasi, Rebecca Mancy, Denise A Marston, Zac Mtema, Malavika Rajeev, Lúcia R. Montebello P Dourado, J. F. Gonzalez Roldan, Kristyna Rysava, Silene Manrique Rocha, Maganga Sambo, Lwitiko Sikana, Marco Vigilato, Victor Del Rio Vilas

## Abstract

**Background:** With a global target set for zero human deaths from dog-mediated rabies by 2030 and some regional programmes close to eliminating canine rabies, there is an urgent need for enhanced surveillance strategies suitable for declaring freedom from disease and elimination of transmission with known confidence.

**Methods:** Using exhaustive contact tracing across settings in Tanzania we generated detailed data on rabies incidence, rabid dog biting behaviour and health-seeking behaviour of bite victims. Using these data we compared case detection of sampling-based and enhanced surveillance methodologies and investigated elimination verification procedures.

**Findings:** We demonstrate that patients presenting to clinics with bite injuries are sensitive sentinels for identifying dog rabies cases. Triage of patients based on bite history criteria and investigation of suspicious incidents can confirm >10% of dog rabies cases and is an affordable approach that will enable validation of disease freedom following two years without case detection. Approaches based on sampling the dog population without using bite-injury follow-up were found to be neither sensitive nor cost-effective.

**Interpretation:** The low prevalence of rabies, and short window in which disease can be detected, preclude sampling-based surveillance. Instead, active case finding guided by bite-patient triage is needed as elimination is approached. Our proposed methodology is affordable, practical and supports the goal of eliminating human rabies deaths by improving administration of lifesaving post-exposure prophylaxis for genuinely exposed but untreated contacts. Moreover, joint investigations by public health and veterinary workers will strengthen intersectoral partnerships and capacity for control of emerging zoonoses.

## Introduction

Dog-mediated rabies kills thousands of people every year in low- and middle-income countries (LMICs)^1^. Yet, with cheap and effective tools available for prevention and control, there are few technical barriers to eliminating rabies as a public health problem, and ultimately to the global elimination of dog-mediated cycles of infection^2^. A target for the global elimination of human deaths from dog-mediated rabies has been set for 2030, with momentum now building in many LMICs^3^. However, clear, pragmatic guidance for effective surveillance to support rabies control and elimination programmes is still lacking.

Surveillance is needed to guide effective management decisions and sensitive surveillance is a prerequisite for verifying pathogen elimination^4,5^. High case detection is required to establish disease absence with certainty, to be confident of interruption of transmission and to rapidly identify introduced cases and secondary transmission^6^. Surveillance has therefore often been tailored during elimination programmes to increase case detection^7^. For example, scarring provided evidence of past smallpox infections^8^; participatory surveillance identified the final outbreaks of rinderpest^9^; and acute flaccid paralysis reports guide surveillance for polio eradication^10^.

Current rabies surveillance guidelines focus on use of validated laboratory diagnostic tests, which are highly sensitive and specific^11,12^ and emphasize the need for notifiability^13^. The World Animal Health Organization (OIE) provisions set out general principles for adequate surveillance systems^14^, but specific recommendations for rabies are limited. In many countries rabies surveillance follows outdated guidance that recommended “a minimum number of samples from suspect cases” be tested, from “between 0.01–0.02% of the estimated population”^15^. Opportunistic or convenience sampling of non-suspicious animals is often conducted to meet such targets e.g. during leishmaniasis surveillance and from culling of “street dogs”. Sample-based targets are widely used to guide veterinary serosurveillance in other diseases, particularly to verify elimination of infection after cessation of vaccination^16^. However, a lack of a consistent or prolonged antibody response to sub-lethal infection precludes this approach for rabies^17^. The challenges of rabies case detection therefore remain substantial: the period during which infection can be detected is short, the infection circulates at low prevalence, and recovering samples from suspected rabid animals is not always feasible^18^. For these reasons, additional and improved approaches to case detection are necessary to guide rabies elimination programmes.

Elimination programmes have potential to reap long-lasting benefits, but, badly managed, they can lead to stakeholder disengagement, programme stagnation and disease resurgence that will be harder to control when political commitment has been lost^19–21^. With efforts underway to control dog-mediated rabies in large parts of the world, and some regions now close to elimination^22^, it is imperative that rigorous and practical verification procedures are developed. Here we use exhaustive contact tracing to generate detailed epidemiological data from populations in Tanzania with differing rabies prevalence. With these data we examine case detection under different surveillance approaches. Specifically, we explore the feasibility, costs and potential for active case finding using bite-patients as sentinels for animal rabies cases in settings in Latin America that are close to eliminating canine rabies.

## Methods

### Study sites

We conducted contact tracing in three sites in Tanzania differing in dog ownership characteristics and rabies prevalence (Figure 1). Serengeti District (2002-2015) is densely populated, mainly agropastoralist, with high levels of dog ownership (i.e. a low human:dog ratio, HDR). Ngorongoro District (2002-2015) is more sparsely populated and inhabited mainly by pastoralists, also with high levels of dog ownership. Pemba Island (2010-2015) is densely populated, with mostly Muslim communities and low levels of dog ownership (Table 1). Control activities have had differing success dependent on effort at each site and geographical isolation, with epidemiological scenarios represented ranging from endemic to local elimination and re-emergence (Table 1).

**Figure 1.**
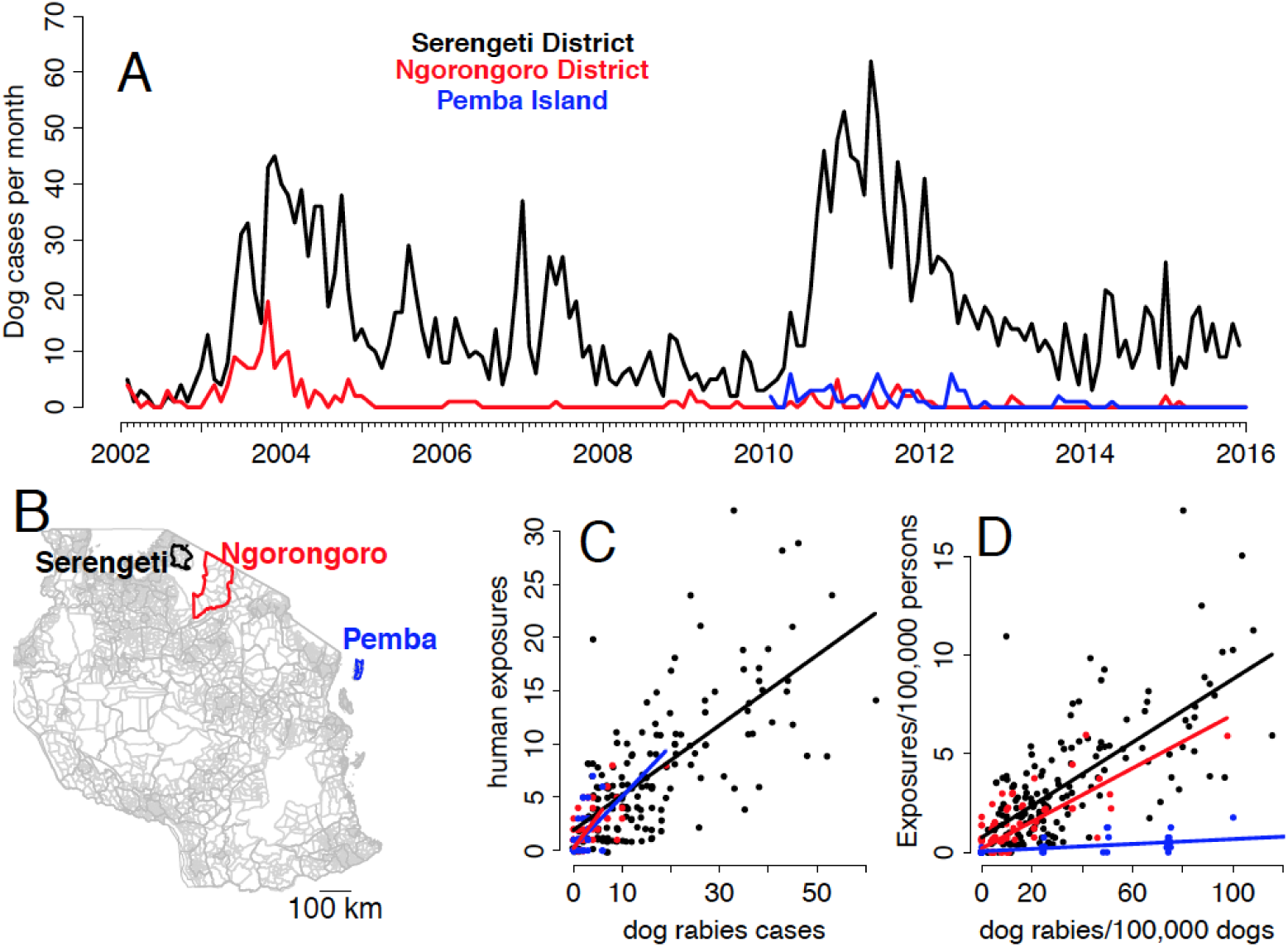
Suspect dog rabies cases and human exposures. A) Time series of suspect rabid dogs in Serengeti District, Ngorongoro District and Pemba Island; B) Study sites in Tanzania, with administrative wards demarcated in light grey and relationships between C) suspect dog rabies cases and human exposures and D) suspect dog rabies incidence and human exposure incidence. Best fitting linear relationships are illustrated for each site.

To compare costs and effort required for current surveillance versus bite-patient triage and investigation of suspicious incidents, we also collected data from areas in Latin America which are close to elimination of dog-mediated rabies (Chiapas state, Mexico and Maranhão state, Brazil). In both countries national dog vaccination programmes have been ongoing for many years and dog rabies cases are now mainly localized to these states. No human deaths from dog-mediated rabies have been reported for several years from either state and rabies incidence in dogs is likely lower than the settings in Tanzania. Surveillance comprises efforts to sample 0.02% of the dog population including opportunistic sampling of dead dogs, and dogs considered suspicious for rabies, as well as quarantining or observation of biting animals.

### Data collection

Contact tracing in Tanzania involved regular collection of health facility records of patients presenting with animal bite-injuries. These records were used as index cases for investigations. We aimed to investigate all incidents as expediently as resources permitted, and always within one year. All bite victims or family members that could be identified from records were interviewed to assess whether the biting animal was rabid using clinical and epidemiological criteria (Box 1). For all ‘possible’ and ‘suspect’ rabies exposures identified, the source of the biting animal and all known persons and animals bitten/ scratched were investigated as well as households the animal was reported to have visited. This iterative procedure was repeated exhaustively for all associated exposures or possible cases identified in animals or humans. Interviews were conducted with assistance of community leaders and veterinary staff, who were encouraged to report other incidents requiring investigation. Wherever possible, brain samples were collected for diagnosis. These were sent in batches for testing to an OIE reference laboratory. Storage in Tanzania and during shipments was not ideal for preservation, therefore samples were tested using a real-time PCR assay^23,24^, as the gold standard OIE tests FAT and RTCIT in particular are sensitive to degradation^25^. More recently samples were also tested in the field using rapid diagnostic tests (RDTs, Bionote, Korea)^26,27^.

Using national census data^28^ and human:dog ratios (Table 1), we estimated projected human and dog populations from 2002 until 2015. Using these denominators, we calculated the incidence of suspect rabid dogs and suspect rabies exposures at these sites.

**Table 1.** Study sites in Tanzania including epidemiological and geographical characteristics and control measures implemented. Human population sizes were from the 2012 National Census^28^. Dog populations were extrapolated from human: dog ratios estimated from household surveys^33–35^ and projected human population sizes. Dog population estimates are shown from 2009.

| Characteristic | Site |  |  |
| --- | --- | --- | --- |
|  | Serengeti | Ngorongoro | Pemba |
| Study Period | 2002-2015 | 2002-2015 | 2010-2015 |
| Human population (2012 census) | 249,420 | 174,278 | 406,848 |
| Dog population (average from 2009) | 50,085 | 22,802 | 3,920 |
| Human: dog ratio | 4.5 | 7 | 100 |
| Dog density/km sq. | 15.3 | 1.5 | 4.1 |
| Mass dog vaccination campaigns | Intermittent since 1990s, with varying coverage | District-wide since 2003, with subsequent lapses | Island-wide campaigns since 2011 |
| Epidemiological scenario | Endemic with resurgences during control lapses | Elimination, re-emergence (2010), now controlled | Eliminated (2014), incursion in August 2016 |
| Geographical Isolation | Bordering high density endemic populations | Bordering low density endemic populations. Rift escarpment acts as barrier | Isolated island. Multiple introductions in recent decades<sup>27</sup> |
| Suspected rabid dogs detected | 2,690 | 184 | 71 |
| Mean annual incidence/ 100,000 dogs (range) | 384 (75-866) | 58 (4-450) | 288 (0-652) |
| Mean annual exposures/ 100,000 persons (range) | 39 (11-99) | 6 (0-36) | 2 (0-7) |

#### Box 1 Case definitions

##### Possible Case

An animal that disappeared, died or was killed within 10 days of showing aggressive behaviour including biting. No reported history of a bite by another animal and no other clinical signs reported.

##### Suspect Case

An animal that disappeared, died or was killed within 10 days of showing any two of the following signs: unprovoked aggression including attempting to bite and grip people, animals or objects; excessive salivation; unexplained dullness/lethargy; hypersexuality; paralysis; abnormal vocalization; restlessness; running without apparent reason. Animal often has a history of a bite. Additional criteria for wild carnivores included loss of fear of humans; diurnal activity of nocturnal species, and unprovoked biting of objects/animals without feeding. No laboratory diagnosis was conducted on human cases, therefore all deaths were identified as suspect rabies from reported symptoms and exposure history.

##### Confirmed Case

Infection confirmed using a high quality diagnostic test (real-time PCR assay) or a rapid diagnostic test.

### Comparison of surveillance approaches

Through simulation, we investigated case detection from surveillance based on sampling targets (0.02% of the overall dog population or of all dead dogs) given levels of suspect rabies incidence in these sites in Tanzania. We estimated average weekly prevalence from detected incidence (Table 1), assuming each rabid dog is symptomatic for one week. To simulate random sampling from the overall population, we drew the target number of samples with equal probability from dog populations equivalent in size to Serengeti, Ngorongoro and Pemba populations, assuming rabies probability for each sample to be equal to estimated weekly prevalence. To simulate sampling of dead dogs, we estimated annual dog deaths in these populations given ***per capita*** mortality of 0.3 per annum^18^. We drew the target number of samples from these estimates of dead dogs, with rabies probability for each sample equal to annual incidence of rabid dogs among dead dogs. For these simulations we assumed perfect diagnostic test specificity and sensitivity. We simulated these experiments 1000 times for each population. From simulation results we calculated the sampling intensity required to detect at least one case with 95% certainty, given observed incidence.

We constructed a probabilistic framework (Figure 2) to estimate case detection from bite-patient investigations (index bite-patients presenting to health facilities only, not subsequent contact tracing, see Appendix), parameterized from contact tracing data from Tanzania. Specifically, we estimated the numbers of persons bitten by suspect rabid dogs; health-seeking behaviour of bite victims; the proportion of bite-patients for which the health status of the biting animal could be evaluated based on clinical history and circumstances of the bite (i.e. rabid/healthy); and the proportion of suspect rabid dogs which disappeared and thus could not realistically have been sampled. Of the samples recovered from suspect rabid animals, we calculated the proportion confirmed positive at WHO/OIE reference laboratories, and using RDTs. To explore the wider applicability of this approach, we investigated differences in suspect rabid dog biting behaviour between sites and the relationship between suspect animal rabies and human exposures using generalized linear mixed models.

**Figure 2.**
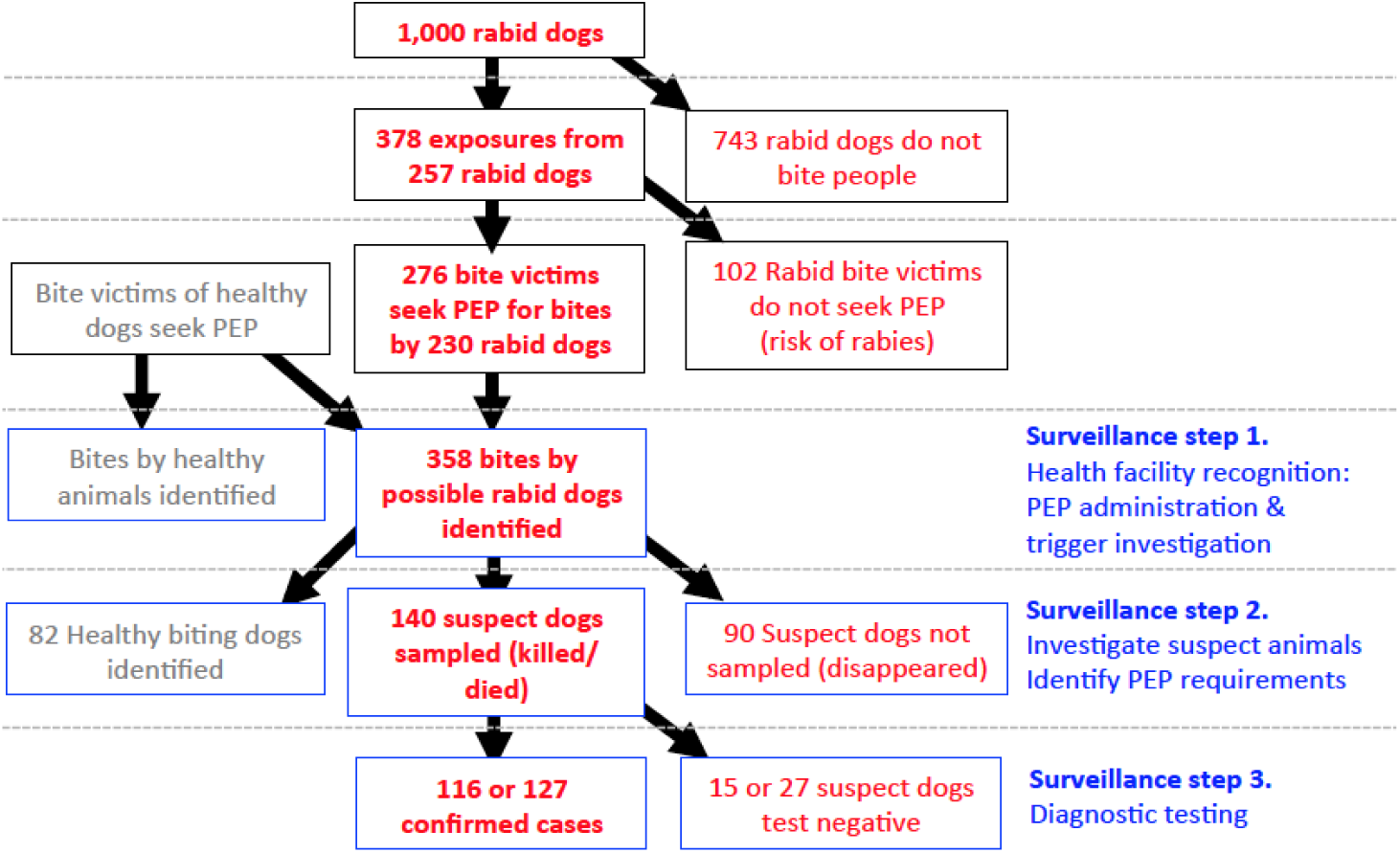
Case detection through active investigation of bite-patients. In this schematic we demonstrate case detection starting from 1000 rabid animals using parameters derived from contact tracing in Tanzania. We assume rabies cases were investigated (red bold), or lost to follow-up (red thin), while exposures from healthy animals were resolved (grey). Surveillance steps are shown in blue. We estimate that 12-13% of circulating rabies cases could be detected in Tanzania.

From stakeholder interviews in Chiapas, Mexico and Maranhão, Brazil, we determined direct costs associated with sampling, laboratory testing and animal observation, including transport, fuel, and equipment associated with investigations, and costs of post-exposure prophylaxis (PEP) for bite-patients. We did not estimate costs of surveillance personnel as these were shared resources conducting a range of other duties.

## Results

### Rabies incidence

From 2,081 investigations of index bite-patients identified from health facility records in Serengeti and Ngorongoro Districts (2002-2015) and on Pemba Island (2010-2015), we identified 1150 suspect rabies exposures. Subsequent contact tracing identified an additional 513 exposures from suspect rabid animals, 3,563 suspect rabid animals (Figure 1) and 64 suspect human rabies deaths, only 32 of whom were recorded in health facilities. Most non-human suspect cases were domestic dogs (83%), followed by livestock (11%), wild carnivores (5%) and domestic cats (2%). Suspect dog rabies incidence varied from zero to 866 cases/ 100,000 dogs/year, with an average of 384 cases/ 100,000 dogs/year in Serengeti, 58 in Ngorongoro and 288 in Pemba (Table 1, Figure 1). Most suspect cases were recorded in Serengeti, the area with the largest dog population (currently >60,000).

### Case detection from sampling-based surveillance

Annually sampling 0.02% of dogs in Serengeti, Ngorongoro and Pemba would require testing 11 dogs, 5 dogs and 1 dog per year respectively. For detected incidence (Table 1) and these sample sizes, the probability of not detecting any cases was >95% given random sampling from these populations or of dying/dead dogs each year (with ∼15,000, 7,000 and 1,200 dog deaths expected in Serengeti, Ngorongoro and Pemba given 0.3 *per capita* annual mortality). Samples required to detect a single rabies case or a minimum of ten cases (for trend evaluation) per year with 95% probability are presented in Table 2. As incidence increases, fewer samples are required to detect rabies, but all sampling-based approaches require impractically large sample sizes (Table 2, Figure S1). Over 40,000 dogs (>80% of the population) need sampling to confirm at least one case with >95% probability in Serengeti, while it would simply not be possible to randomly sample enough dogs in Ngorongoro or Pemba to be confident of confirming rabies because at times there may not be any infectious dogs in the population from which rabies virus could be detected (Figure S1). Focusing on dead or dying dogs requires over 230 samples (1.6% of all deaths) in Serengeti to confirm at least one case with >95% probability, over 1,500 samples (>20% of deaths) in Ngorongoro and 300 in Pemba (>25% of deaths).

### Case detection from active investigations of bite-patients

During contact tracing, sufficient details about the circumstances and animal history were recalled to differentiate suspect rabid from healthy biting animals in most incidents (98%, Figure 2, Table 3). The majority of patients reported due to exposures from suspect rabid animals (58%). Of the remaining bite-patients, 8% were by animals of ambiguous status that could not be resolved, and the rest were due to healthy animals; many of these patients lived near the health facilities (data not shown). From contact tracing, we quantified the proportion of victims of suspect rabid bites that sought health care (Figure 2). Over >25% of suspect exposures did not attend a clinic or obtain PEP, with lowest attendance in Serengeti, and higher attendance in Ngorongoro and Pemba (Table S1). Many bite victims on Pemba identified through contact tracing provided a receipt of PEP, but could not be identified in official records, indicating poor recording.

After exhaustive contact tracing, we identified a total of 2944 rabid dogs from 1113 suspect human exposures i.e. 26% of suspect rabid dogs were traced to a human exposure. Assuming that these numbers are an adequate estimate of total infections and exposures, we estimate a mean of 0.38 persons exposed per suspect rabid dog (Figures 2 and 3). Rabid dog biting behaviour was variable, with extreme examples of one suspected rabid dog that bit 18 people and another that bit >70 dogs (several of these bitten animals were subsequently confirmed rabid). We did not detect significant differences in dog biting behaviour between sites despite major differences in dog density (and human:dog ratios). At each site, numbers of people bitten by rabid dogs each month were similarly correlated with numbers of suspect rabid dogs, but correlations between monthly exposure incidence and suspect dog rabies incidence were markedly different between sites, i.e. the average number of persons bitten per rabid dog was similar across communities, and bites per person was higher in communities with more dogs, (Figure 1C,D).

**Figure 3.**
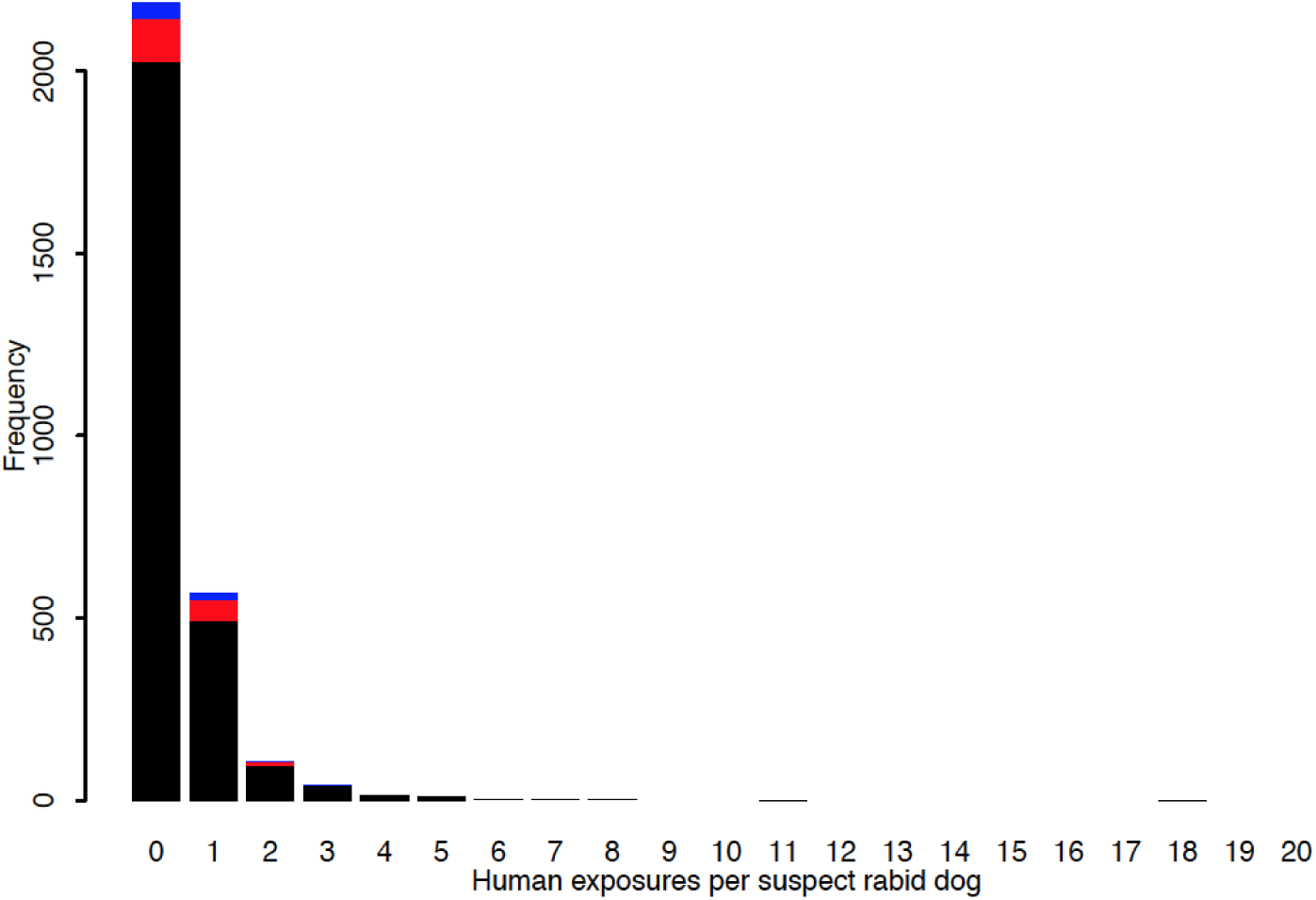
Individual variation in biting behaviour of rabid dogs in Serengeti district (black), Ngorongoro district (red) and Pemba Island (blue).

We estimated that diagnostic samples could be recovered from ∼61% of suspected rabid animals if promptly investigated (Figure 2); 15% of suspect dogs died (452), 46% were killed (1352), while the remaining 39% were reported to have disappeared (1150) therefore presumably could not have been sampled. The potential probability of sample recovery was higher in Ngorongoro than Serengeti or Pemba because rabid dogs were more likely to be killed here (60% versus 45% and 44% in Serengeti and Pemba). In practice, we obtained samples from 13% of suspected animal rabies cases identified from contact tracing. Of these samples, 83% of those sent for laboratory testing, and 90% of those tested using RDTs were confirmed positive (Tables 2 and 3).

Accounting for multiple biting by rabid dogs, we estimate that in Tanzania ∼23% of dog rabies cases can be identified through bite-patient triage (Figure 3, Table 3), without further contact tracing. If a higher proportion of persons bitten reported to clinics case detection would be higher, with up to 26% of rabid dogs identified if all rabies-exposed bite victims sought care. Assuming proficient laboratories or reliable RDTs, we estimate that subsequent investigations could confirm between 12% to 14% of cases (Figure 3, Tables 2 and 3).

### Surveillance costs

In Chiapas, 578 bite-patients/year presented to facilities from 2011 to 2014 (11 bites /100,000 persons/year). From an average of 101 dog samples submitted annually only 6 (1.5%) tested positive – all from municipalities with high bite incidence. In Maranhão, 8,541 bite-patients attended clinics annually between 2010 and 2014 (125 bites/100,000 persons/year). A higher proportion of samples tested positive (15%, 212/1,417), with a correlation between testing and positivity (p<0.001). This indicates that a proportion of sampled animals were targeted because of being suspicious for rabies and were not randomly sampled. In contrast, animals culled for leishmaniasis surveillance in Maranhão all tested negative for rabies (Figure S2).

Estimated costs of investigating bite-patients in Chiapas and Maranhão were similar to current surveillance (we expect costs to be closest to the lower end of the ranges reported in Table 4). Following triage, only a small proportion of patients would likely require investigation, therefore enhanced surveillance could be conducted by current surveillance personnel at no extra cost. Costs would be expected for keeping dogs under observation given ambiguity in identifying rabies from interviews (Table 4), while PEP costs could be reduced with identification of bites by healthy animals.

**Table 2.** Case detection from sampling-based surveillance versus investigations of bite-patients. Case detection was estimated as the percentage of all cases detected using the specified surveillance strategy. We assumed a dog population size equivalent to that of an average state in Mexico (˜409,000 dogs), and compared case detection under high and low incidence settings (384 vs 58 rabid dogs/100,000 dogs/year, similar to Serengeti or Ngorongoro district respectively). For bite-patient investigations we assumed bite incidence equivalent to that recorded for Chiapas state, Mexico with the same proportion of ambiguous bites as in Tanzanian settings. Not all ambiguous bites would need investigation to confirm rabies circulation, but all would need investigation to verify disease freedom.

| Surveillance strategy | Target | Incidence (per 100,000 dogs) | Average case detection (numbers positive) | Sampling to detect 1 case (10 cases) |
| --- | --- | --- | --- | --- |
| Sampling-based | 0.02% of population | High (383) | 0.0004% (0.006) | >40,000 (>200,000) |
|  |  | Low (58) | 0.0004% (0.0009) | >270,000 (NA) |
| Sampling-based | 0.02% dead/dying dogs | High (383) | 0.02% (0.3) | >230 (>1,200) |
|  |  | Low (58) | 0.02% (0.05) | >1500 (>8,000) |
| Bite-patient sentinels | Rapid diagnostic testing | High (383) | 13-14% (48-54) | <50 (<67) probable bite-patients |
|  |  | Low (58) | 13-14% (7-8) | <50 (<67) probable bite-patients |
| Bite-patient sentinels | Laboratory testing | High (383) | 12-13% (44-50) | <50 (<67) possible bite-patients |
|  |  | Low (58) | 12-13% (7) | <50 (<67) possible bite-patients |

**Table 3.** Probability steps for estimating case detection based on investigations of bite-patients. Parameters derived from contact tracing in Tanzania, based on schematic in Figure 2. Case detection is based on health-seeking behaviour observed in Tanzania and assuming all bite victims seek treatment (maximum). Estimates exclude attendance by victims bitten by other species (domestic cats, wildlife and livestock) which would increase case detection. *identification of patients bitten by possible rabid animals is included (see Figure 2), but does not affect case detection, only the number of investigations required (Table 2).

| Step | Probability | N |
| --- | --- | --- |
| A rabid dog biting at least one person | 0.26 (0.24-0.27) | 2,944 |
| At least one person bitten by rabid dog attends clinic | 0.89 (0.86-0.91) | 758 |
| Follow up with bite patient identifies biting dog status | 0.98* (0.97-0.98) | 3044 |
| Rabid dog can be sampled (killed/died) | 0.61 (0.59-0.63) | 2945 |
| Positive sample using rapid diagnostic test | 0.90 (0.85-0.94) | 175 |
| Laboratory confirmation of suspicious sample | 0.83 (0.78-0.87) | 313 |
| Case detection based on clinical suspicion | 0.23 (max: 0.26) |  |
| Case detecting based on RDT | 0.13 (max: 0.14) |  |
| Case detection based on laboratory confirmation | 0.12 (max: 0.13) |  |

**Table 4.** Estimated annual costs of current versus enhanced surveillance in Maranhão, Brazil and Chiapas, Mexico. Brazil population data is from 2012:www.ibge.gov.br/home/estatistica/populacao/estimativa2012/estimativadou.shtm. Mexico population data is from 2015: www.inegi.org.mx/est/contenidos/proyectos/encuestas/hogares/especiales/ei2015/doc/eic2015presentacion.pdf). *In Maranhão, costs exclude laboratory testing which is carried out under centralized state funding. The range of costs for bite investigations was calculated under low and high incidence (58-383 rabid dogs/ 100,000 per annum as in Ngorongoro and Serengeti, Tanzania). However, as high incidence resulted in more bites than were recorded from Chiapas, we assumed that all bites were due to rabid animals, which is unlikely. We assumed that 8% of bites were ambiguous and required observations (8% of bites in Tanzania that were not due to suspect rabid dogs, were ambiguous). PEP costs were based on average costs per exposure.

| State | Population | Activity | Current costs (USD) | Costs of bite-patient investigations (USD) |
| --- | --- | --- | --- | --- |
| Chiapas, Mexico | 5,217,908 | Sampling & testing | 4,881 | 3,274 - 17,112 |
|  |  | Observation of dogs | 7,620 | 2,312 |
|  |  | PEP | 15,510 |  |
| Maranhao, Brazil | 6,714,314 | Sampling & testing* | 10,068 | 3,943 - 26,247 |
|  |  | Observation of dogs | NA | 33,625 |
|  |  | PEP | 189,804 |  |

## Discussion

With the 2030 global target for zero human deaths from dog-mediated rabies approaching^3^, practical and affordable surveillance criteria to guide elimination programmes are urgently needed. We propose that using bite victims as sentinels for animal rabies is a targeted way to increase case detection that could have great utility as rabies is increasingly controlled. We found that investigating bite-patients can detect and confirm >10% of dog rabies cases in challenging settings at low cost. From previous work demonstrating that 5% case detection is sufficient to verify rabies freedom^30^, we recommend that, by implementing surveillance systems based on investigations of bite-patients, elimination of dog rabies could be validated in areas that detect no cases for two or more years.

We demonstrate that sampling-based surveillance is neither sensitive nor cost-effective for rabies and does not provide statistically robust evidence on the occurrence of rabies in populations, irrespective of incidence levels. Moreover, the implications of this approach has included opportunistic, costly and unnecessary killing and testing of animals not suspect for rabies. At times guidelines have been misinterpreted with authorities targeting 0.2% or even 2% of the population with ramifications for dog welfare and conflict between authorities and communities.

We predict that investigations of bite-patients will be a robust approach for rabies surveillance that can be applied across settings. We did not detect substantive differences in rabid dog biting behaviour among Tanzanian populations, which might affect its applicability. However, we do expect differences in human behaviour, both in terms of health seeking and actions taken against rabid dogs. The higher the proportion of people seeking PEP in the event of a bite from a suspect rabid dog, the higher proportion of cases detected. However, even in Tanzania, where a relatively high proportion of bite victims do not seek treatment (27%, Figure 3), due to lack of awareness, financial barriers or limited PEP availability^31^, this approach is still practical. In wealthier countries with advanced rabies control programmes we expect this proportion to be lower and therefore case detection to be higher. More victims of bites by healthy animals are also likely to present to health facilities, particularly where PEP is more accessible (most bite victims in Tanzania pay >$10/dose^32^), introducing costs associated with investigations. However, questions administered by health workers should identify both patients requiring investigation, and patients for whom PEP is unnecessary or could be discontinued. Judicious PEP use reduces costs while resulting investigations of suspect animals could reveal victims in need of PEP who did not seek care. Indeed, an action-orientated approach to surveillance^33^ such as investigation of bite-patients, measures progress towards the Sustainable Development Goals (Indicator 3.3.5, the number of people requiring interventions against Neglected Tropical Diseases, which includes PEP). We suggest a protocol for this enhanced surveillance approach in the Supplementary Material.

Timeliness of investigations will influence the effectiveness of using bite-patients as sentinels. Delays in investigations will reduce the number of samples that can be recovered, may compromise prompt PEP administration and will delay detection and therefore any outbreak responses. Widespread network coverage and phone ownership could facilitate timely follow up to identify suspect dogs requiring investigation, including observation/quarantine if alive. Joint investigations are an ideal way to strengthen inter-sectoral relationships between health and veterinary sectors, urgently needed to investigate and respond to emerging zoonoses.

Investigations of bite sentinels does not negate the importance of on-going investigations of suspect animal cases, including those that are not linked with human exposures. Bite-patient investigations are a complementary tool to improve recovery of samples from suspicious animals including those diagnosed when sick or dead animals brought directly to veterinary clinics or disease control centres. Any potentially rabid animal, irrespective of whether they bit anyone, should be treated carefully and investigated with urgency. Indeed bite-patient triage and investigation of patients bitten by suspect animals should detect other rabies variants and guide PEP use.

Effective surveillance is integral to disease control and elimination. As countries progress towards rabies elimination, their first step will be to build a dossier of epidemiological evidence for the disease status in their region, interventions that have led to this status and surveillance and response capacity to prevent reestablishment. A key aspect of this dossier is adequate and representative surveillance. We recommend triage of bite-patients and investigation of suspicious cases to enhance surveillance to validate elimination of dog-mediated rabies and maintain freedom through early detection and responses to incursions.

## Authors contributions

KH designed the study, collected data, analysed the data and wrote the first draft. KB, KR, KHo, MR, MC collected data and contributed to interpretation. JC, AL, KL, ZM, MS, LS, STMB, SMR, EC, VG, LRMPD, JFGR, MV collected data. ARF and DAM contributed reagents and diagnostic capability for confirming rabies and edited the manuscript. JD, BAR, DTH, SC, RB and VDRV edited the manuscript.

## Ethics Committee Approval

This work was approved by the Institutional Review Board of Ifakara Health Institute and the Medical Research Coordinating Committee of the National Institute for Medical Research of Tanzania (NIMR/HQ/R.8a/Vol.IX/946).

## Acknowledgments

We are grateful to Tanzania Ministries of Health and Social Welfare, and of Agriculture, Livestock and Fisheries, the Zanzibar Ministry of Health and Research Council, the WHO Country Office-Tanzania, the National Institute for Medical Research and the Tanzanian Commission for Science and Technology for permissions and support. We thank rabies control programme staff, district veterinary officers, livestock field officers, health workers and community members who assisted with this study in Tanzania, as well as rabies control staff from the Secretary of Health of the state of Maranhão in Brazil and from the Ministry of Health in Mexico.

Funding was provided by the Wellcome Trust (095787/Z/11/Z to KH, 103270/Z/13/Z to KL), the UBS Optimus Foundation, the Pan American Health Organization, the Medical Research Council and the Research and Policy for Infectious Disease Dynamics Program of the Science and Technology Directorate, Department of Homeland Security, Fogarty International Centre, National Institute of Health. Dog vaccinations in Southern Tanzania were supported by the Bill and Melinda Gates Foundation, and vaccines were donated by MSD Animal Health to campaigns in Northern Tanzania.

## Research in Context

### Evidence before this study

We searched PubMed for studies published in any language up until December 2016 on “rabies” and “surveillance” and “case detection” or “elimination”. Almost all studies focused on methods of laboratory diagnosis, with none investigating means of increasing identification of potential case and recovery of samples for subsequent laboratory confirmation. Many studies alluded to underreporting and under detection of rabies, and a much higher burden of disease than confirmed cases suggest. A previous modelling study highlighted the need for improved case detection to validate elimination of transmission and more generally to improve responses to rabies outbreaks. Otherwise there was an absence of scientific recommendations for practical surveillance strategies, which are urgently needed to inform international guidelines for rabies control and elimination programmes.

### Added value of this study

In this study we used detailed and comprehensive contact tracing data from Tanzania to measure rabies incidence in different settings. Using these data we showed that detection methods based on sampling a proportion of the dog population do not provide useful guidance for rabies control strategies, whereas an approach focusing on bite-patients as targeted sentinels can lead to detection and laboratory confirmation of over 10% of animal rabies cases.

### Implications of all the available evidence

Our study suggests that investigations of suspicious incidents, following triage of bite-patients using clinical criteria and bite history, are an affordable and effective strategy to improve case detection. Together with evidence from a previous modelling study, we conclude that using this methodology, it should be possible to validate rabies freedom given two years without detection of rabies cases. Moreover, the improved detection of rabies with the implementation of bite-patient investigations should also improve the administration of lifesaving human post-exposure vaccines and support rapid and more effective responses to incursions. We therefore suggest the use of bite-patient triage and investigation of suspicious incidents to improve case detection in settings around the world that are now close to eliminating canine rabies.

## Supplementary Material

### Appendix: Recommended protocol for enhanced surveillance to guide rabies elimination programmes and verify freedom from disease

1. Contact information should be recorded for all patients reporting to health facilities due to animal bites or scratches, and a triage process conducted to determine PEP requirements and whether further investigation is required.
2. Health workers should ascertain whether the biting animal was possibly rabid, showing at least two of the following signs:

a. unprovoked aggression including attempting to bite and grip people, animals or objects
b. excessive salivation
c. unexplained dullness/lethargy
d. hypersexuality
e. paralysis
f. abnormal vocalization
g. restlessness
h. running without apparent reason
i. loss of fear of humans (for wild animals)
j. diurnal activity of nocturnal species (for wild animals)
k. unprovoked biting of objects/animals without feeding (for wild animals)
3. If the animal showed any of the aforementioned signs, an investigation should be conducted and PEP initiated. Investigations should be coordinated between health and veterinary staff and the extent/type of investigation should depend on the outcome of the biting animal as follows:

a. Animals that disappeared after the bite should be considered suspect for rabies, but lost to follow up (no sample for laboratory diagnosis). PEP should be continued as a precautionary measure. Other indicative signs of rabies include the animals being of unknown origin or having a history of a bite by another (possibly rabid) animal. Interviews with the dog owner/ family, if known, should be conducted to identify other persons in need of PEP.
b. For animals that died or were killed after the bite, samples should be collected and tested and PEP continued as a precautionary measure (irrespective of the test outcome).
c. Animals alive at the time of patient presentation to the health facility should be observed or quarantined until 10 days after the date of the bite. Phone call follow up could be used to confirm whether the dog is alive at the end of this period or to notify healthy/veterinary staff if the animal becomes sick or dies.

i. If the animal dies during this period the patient should complete PEP and a sample should be collected and tested.
ii. If the animal shows further signs of rabies, the patient should complete PEP and the animal should be euthanized and samples collected for diagnostic testing
iii. If the animal is alive and healthy at the end of this period, PEP can be safely discontinued.

### Surveillance should record

1. the number of patients presenting to clinics with animal bite injuries
2. the number of patients requiring investigation due to potential exposure from possible rabid animals
3. the outcome of investigations, specifically:

a. numbers of negative animals – alive and healthy after ten day period
b. number of animals lost to follow up (animal disappeared, sample could not be recovered), therefore considered suspect for rabies
c. number of confirmed rabies cases (positive laboratory diagnosis)
d. number of indeterminate cases (suspicious for rabies but negative/ indeterminate test result)

Validation of freedom should be based on review of these numbers (1-3 inclusive) recorded monthly over a period of at least two years. If no cases of canine rabies are confirmed over a two-year period, an area could be certified free from rabies. However, evidence that investigations were conducted should be presented (up to 8% of bite patients are expected to be investigated) and of laboratory proficiency for confirming cases. It is expected that around 40% of suspect biting animals will be lost to follow up (disappeared after the bite). If a significantly larger proportion of biting animals are lost to follow up, further monitoring should be undertaken. For example, if highly suspicious rabies cases are detected on the basis of clinical criteria, freedom should not be declared until two years after the suspect clinical case with no further laboratory confirmation of cases.

### Examples of triage process and surveillance investigations

### 1. *Triage identified bite patient requiring investigation - euthanasia of suspect dog and case confirmation*

Margaret (8y, female) was bitten by the family puppy. The family did not initially seek hospital care until 5 days later, when the puppy began biting objects, snapping at flies and barking excessively. The health worker interviewed Margaret and Margaret’s mother who brought Margaret to the clinic. They identified that the puppy had abnormal vocalization, was restless, and showing unprovoked aggression. The health worker contacted the local veterinary officer who visited Margaret’s house that afternoon. Margaret’s father explained to the veterinary officer that he had adopted the puppy, which he had found in the next village. The puppy had been tied to the gate, but was continuing to behave strangely and was now trying to bite everything that came near, so the veterinary officer euthanized it and collected a brain sample. The sample tested positive using a rapid diagnostic test. Margaret’s brother was also sent to get PEP when he revealed that the puppy had scratched him while he was trying to restrain it. Both Margaret and her brother completed PEP.

### 2. *Triage identified bite patient requiring investigation – quarantine of suspect dog and subsequent case confirmation*

Chacha (6y, male) was bitten on his calf by the neighbour’s dog while he was going through the neighbour’s gate. It was a severe bite that was bleeding a lot. His parents took him to the clinic to take care of the wound. Although the health worker determined that the dog was behaving aggressively when it bit Chacha, no other signs of rabies were evident. Following coordination with the health worker, the local veterinary officer visited Chacha’s family and the neighbour who owned the biting dog, to conduct an investigation. The neighbour reported that the dog had been bitten a few weeks previously by an unknown dog during the night that had subsequently disappeared. On observing the dog that bit Chacha, it was clear that the animal was biting objects and barking excessively without provocation. The veterinary officer took the dog to the local veterinary office for quarantining, however the dog died on the second day in quarantine. A brain sample was collected and tested positive for rabies under the FAT. Chacha completed the course of PEP.

### 3. *Triage identified bite patient requiring investigation - sample collection and case confirmation*

Sara (16y, female) was bitten by an unknown cat while she went outside to the toilet in the night. The cat gripped Sara’s leg without letting go. She called for help and her mother came and strangled the cat while it continued to grip Sara’s leg. The next morning Sara and her mother attended the local clinic. The health worker started both Sara and her mother on a course of PEP (Sara’s mother was also scratched by the cat during this encounter) and contacted the local veterinary officer. The veterinary officer went straight to Sara’s house and collected a sample from the cat, which had been thrown into the rubbish pit. The sample tested positive for rabies.

### 4. *Triage identified bite patient requiring investigation - suspect case lost to follow up (no sample collected or tested)*

Teresa (55y female) was bitten on the ankle by a dog that came to her house. The health worker initiated PEP for Teresa and contacted the local veterinary officer to conduct an investigation. The veterinary officer visited Teresa’s house and interviewed family members who confirmed that the dog was of unknown origin. One of Teresa’s children thought that the same dog had also bitten a child from the neighbouring house. The veterinary officer visited the neighbour to find out more information and confirmed that the same dog had also bitten a 9-year old boy before it was chased away. The veterinary officer sent the boy to the local clinic for PEP and advised both households and the local village leader to report any sightings of the suspicious animal immediately. Both Teresa and the boy completed PEP, but no sample was ever obtained from the animal, which was not seen again.

### 5. *Triage identified bite patient as bitten by a healthy animal - PEP initiated but discontinued on subsequent phone call follow up*

Simon (10y, male) was bitten by the family dog when arriving home from school. The next day, his parents took him to the local health facility, where he was started on a course of PEP. However no other signs of rabies were identified by the health worker, therefore he advised the family was advised to watch the dog and call the health facility and local veterinary officer if the dog started to behave strangely over the next week. After ten days, the health worker called to find out whether the family dog that bit Simon was still alive. Simon’s parents confirmed that the animal was well, and the health officer advised them that they were not required to return to the clinic for the rest of the PEP course.

### 6. *Triage identified bite patient requiring investigation - confirmation of a healthy animal and discontinuation of PEP*

Juma (20y, male) attended the local hospital after his neighbour’s dog bit him. He did not know whether the dog was still alive, and so the health worker initiated an investigation. The local veterinary officer visited Juma and his neighbour to find out more about the status of the dog. The dog had been vaccinated earlier that year and had a litter of puppies less than one week old. The neighbour confirmed that Juma had been bitten by the dog when he had approached the shed where the puppies were. The local veterinary officer advised that Juma discontinue the course of PEP and contacted the health worker to confirm that the biting dog was healthy and not rabid.

### 7. *Triage identified bite patient bitten by a healthy animal - PEP not initiated*

Tina (65y, female) stepped on the tail of her pet dog, which then bit her on the hand. After a few days the wound became infected and so she attended the local health facility, where the wound was cleaned and she was given antibiotics. The health worker was able to determine from Tina that the dog was alive and healthy and previously vaccinated, and therefore she was not advised to start PEP.

**Figure S1.**
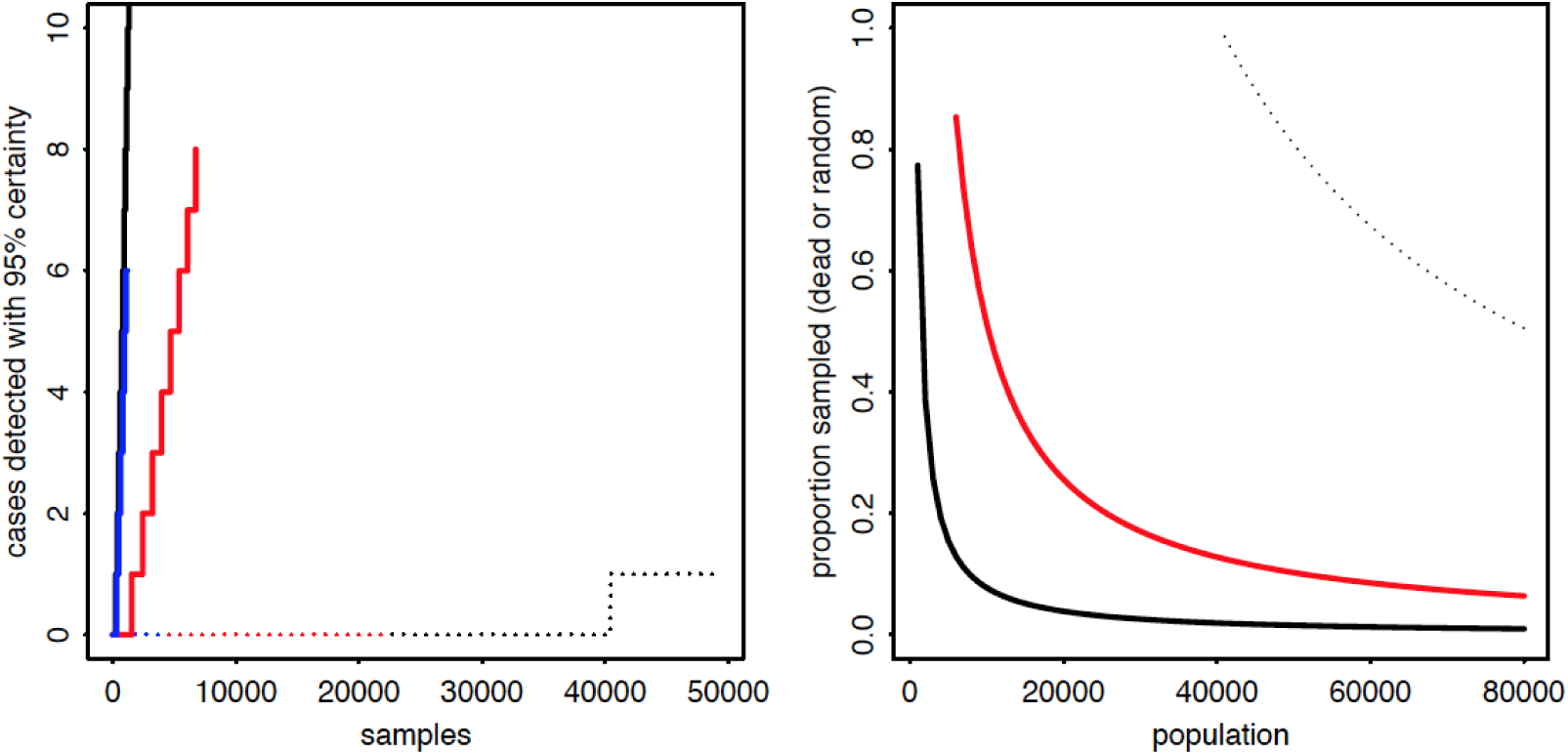
Power to detect rabies cases with 95% certainty in A) Serengeti district (black), Ngorongoro district (red) and Pemba Island (blue) according to their incidence, based on random sampling from entire population (dotted lines) versus random sampling of dead/dying dogs (solid lines) and B) the proportion of the population that must be sampled to detect at least one case in high versus low incidence populations (black: ∼380 cases/100,000 dogs; red: ∼60 cases/100,000 dogs), based on random sampling (thin dotted lines - not possible to detect in low incidence populations) or as a proportion of dog deaths (solid lines - asymptoting at 230 dead dogs in high incidence populations and 1500 in low incidence populations).

**Figure S2.**
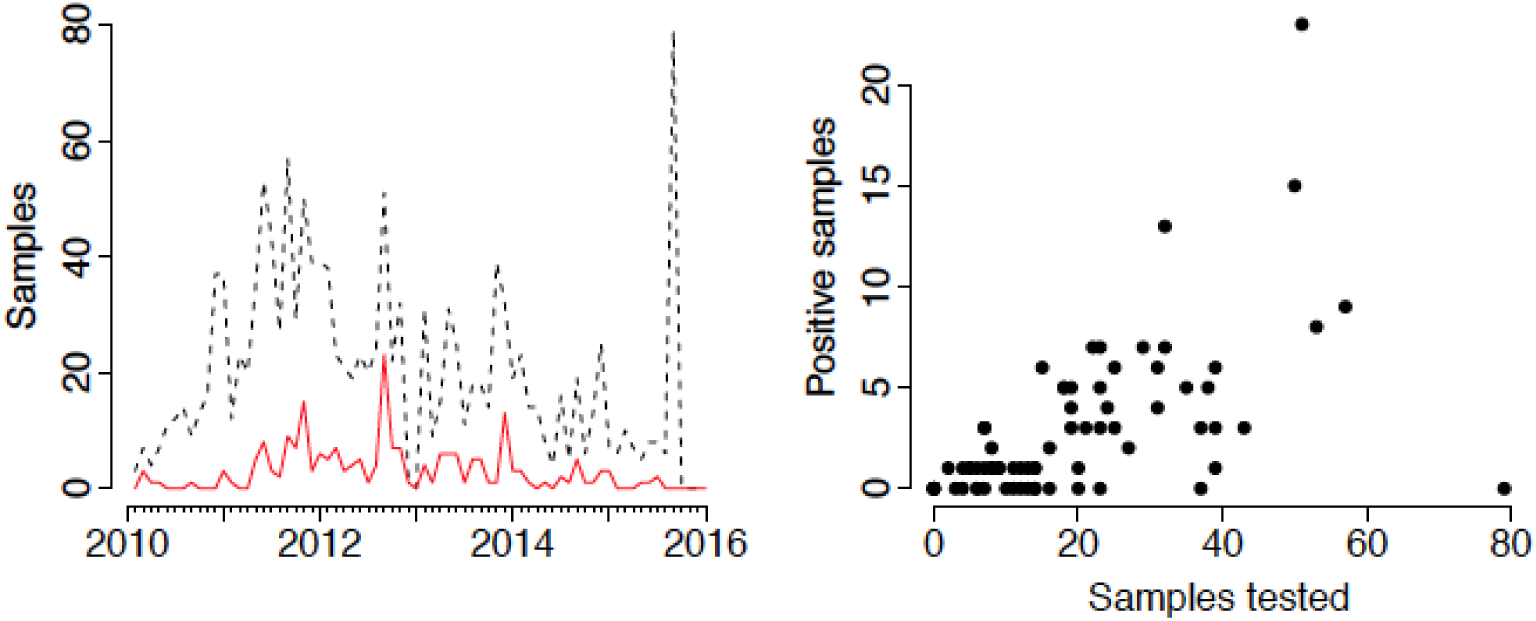
Relationship between samples submitted and tested positive in Maranhão, Brazil. A) Time series of samples tested (dashed black line) indicating which samples tested positive (solid red line) and B) relationship between number of samples tested per month and samples that tested positive. There is a strong correlation between samples tested and percentage positive, although an outlier is evident from one month when a large number of samples were tested from dogs that were culled for leishmaniasis surveillance (n=79).

